# Visual search efficiency is modulated by symmetry type and texture regularity

**DOI:** 10.1101/2024.06.24.600439

**Authors:** Rachel Moreau, Nihan Alp, Alasdair DF Clarke, Erez Freud, Peter J. Kohler

## Abstract

More than a century of vision research has identified symmetry as a fundamental cue, which aids the visual system in making inferences about objects and surfaces in natural scenes. Most studies have focused on one type of symmetry, reflection, presented at a single image location. However, the visual system responds strongly to other types of symmetries, and to symmetries that are repeated across the image plane to form textures. Here we use a visual search paradigm with arrays of repeating lattices that contained either reflection or rotation symmetries but were otherwise matched. Participants were asked to report the presence of a target tile without symmetry. When lattices tile the plane without gaps, they form regular textures. We manipulated texture regularity by introducing jittered gaps between lattices. This paradigm lets us investigate the effect of symmetry type and texture regularity on visual search efficiency. Based on previous findings suggesting an advantage for reflection in visual processing, we hypothesized that search would be more efficient for reflection than rotation. We further hypothesized that regular textures would be processed more efficiently. We found independent effects of symmetry type and regularity on search efficiency that confirmed both hypotheses: visual search was more efficient for textures with reflection symmetry and more efficient for regular textures. This provides additional support for the perceptual advantage of reflection in the context of visual search and provides important new evidence in favor of visual mechanisms specialized for processing symmetries in regular textures.

## 1. Introduction

As we move through the world, the brain generates our visual experience by rapidly processing a constant stream of visual stimuli. Despite the apparent effortlessness of vision, this process is highly complex. Seminal perception research proposed that visual processing is simplified through a set of fundamental gestalts which provide structural limitations on the interpretation of visual stimuli (Wertheimer, 1923). In the current study, we investigate how one of these fundamental gestalts; symmetry, contributes to the perception of textures and how this impacts the efficiency of visual processing.

Symmetries are prevalent in the natural world and can be found in man-made objects throughout human history (Jablan, 2002). Multiple studies have identified a key role for symmetry in scene and object perception (Bertamini, Silvanto, Norcia, Makin & Wagemans, 2018), contributing to behaviors as fundamental as shape perception (Bahnsen, 1928; Machilsen et al., 2009) and as sophisticated as judgments about facial attractiveness (Grammer and Thornhill; 1994). Much of this literature has focused on reflection or mirror symmetry, but reflection is only one of four fundamental symmetry types, with the others being: rotation, translation, and glide reflection. While reflection can be seen in the bilateral bodies of many animals and is especially behaviorally relevant for human faces, there are examples of every symmetry type in nature, e.g., rotation symmetry in flower petals, honeycombs, and snowflakes.

This raises the question: how does the visual system process these various symmetry types, and how do they differ from previous findings with reflection symmetry? Since the beginning of symmetry as a research topic in vision research, reflection has been considered unique among the symmetry types (Mach, 1897, eng. translation 1959). Psychophysical studies show that reflection symmetry can be detected pre-attentively, requires less cognitive resources, and allows for faster reaction time than rotation and translation (Wagemans 1995, Wagemans 1997, Olivers & Van Der Helm, 1998; Treder, 2010; Bertamini & Makin, 2014). It has been suggested that the advantage of reflection might be a result of evolutionary pressures to optimize encoding of behaviorally relevant stimuli that have reflection symmetry, such as faces (Grammer & Thornhill, 1994).

Most studies on the role of symmetry in visual behavior have considered one or two axes of symmetry centered on a single location in the image, consistent with the way symmetries would most likely occur over objects in the natural world (Bertamini, Silvanto, Norcia, Makin & Wagemans, 2018). However, symmetries can also be found in regular textures known as the wallpaper groups – a set of 17 unique combinations of the four fundamental symmetry types (Fedorov, 1891; Polya, 1924; Liu et al., 2010). The wallpaper name is apt as the textures resemble a Victorian wallpaper or rug (see Figure 1). Regular and near-regular textures are abundant in natural and man-made environments (Liu, Lin & Hayes, 2004), and symmetries in regular textures generate strong responses in the visual cortex of humans (Kohler et al., 2016; Kohler et al., 2018; Kohler & Clarke, 2021; Alp et al. 2018) and other primates (Audurier et al., 2021).

**Figure 1:**
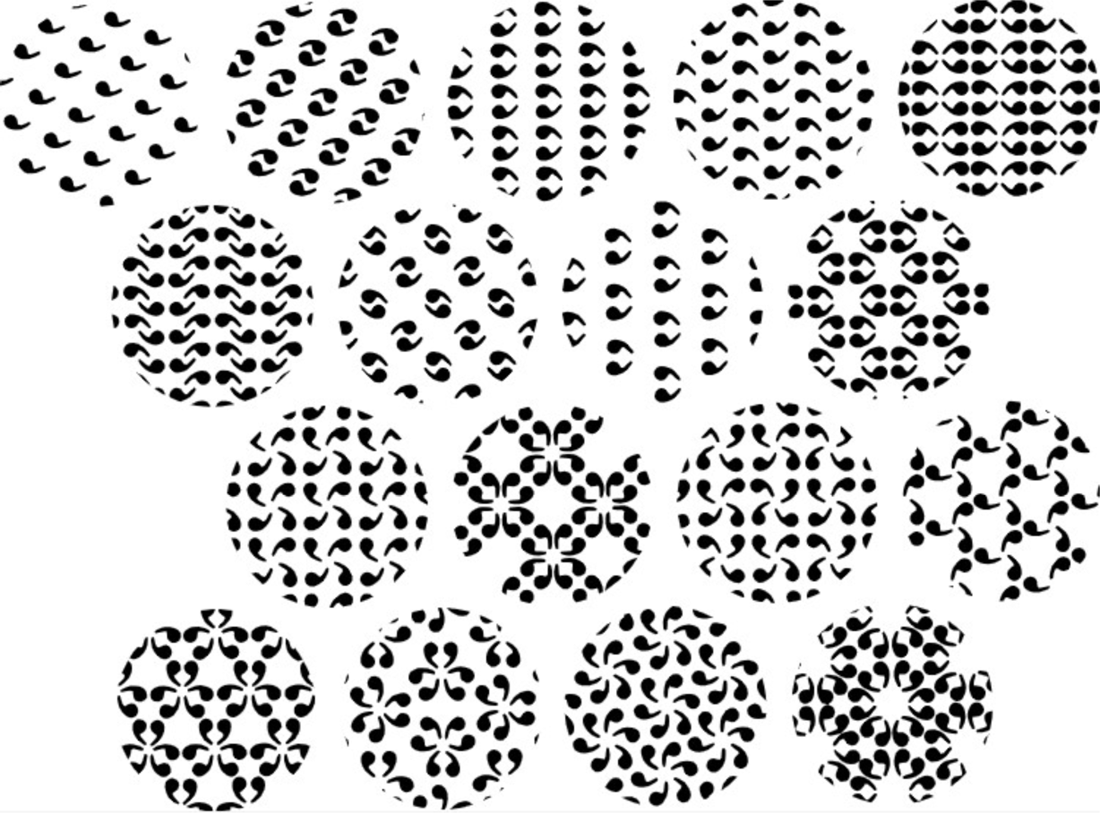
The 17 wallpapers rendered with a comma-like symbol as the repeating element. Illustration based on Wade (1993).

The growing literature on wallpaper groups shows that when embedded in regular textures, each of the different symmetry types can give rise to reliable responses in the visual cortex. The behavioral consequences of this have yet to be explored. The current study seeks to address that gap in the literature by investigating the efficiency of processing reflection and rotation symmetries when these symmetries are presented in regular textures and when they are not. This will provide valuable information about how the human visual system handles complex representations of symmetry and regularity, and how these striking patterns may contribute to the perception of natural scenes.

We addressed these questions using a visual search task. Visual search has been used to probe the extent to which a given cognitive process takes place in a serial or parallel fashion. Reaction time and accuracy are measured as participants search for a target. Typically, the target is either presented among distractors or hidden in noise. If information is being processed serially, the observer has to scan through each individual array element until the target is found. Thus, adding more distractors to the array will result in longer reaction times and/or lower accuracy. On the other hand, if information is processed in parallel, the target will “pop-out”, resulting in reaction times and accuracy that are constant across array sizes (Treisman & Gelade, 1980). Previous studies have utilized visual search to dissociate parallel or serial processing of visual properties such as colour and orientation. It has been found to be an effective way of differentiating visual properties based on the cognitive resources required for processing (Bundesen, Kyllingsbæk, Larsen, 2003; Kyllingsbæk & Bundesen, 2007; Cavanagh, Arguin, Treisman, 1990). Serial and parallel processing are often presented in binary fashion, as two qualitatively distinct types of cognitive processing. However, it is likely more realistic to conceptualize them as endpoints along a spectrum (Wolfe, 1998, 2016). That is, cognitive processes may operate as strictly serial or parallel, but can also fall anywhere between the two. In the current study, we therefore compare conditions in terms of “more serial” or “more parallel”.

Visual search has been used to study symmetry in two main ways: through within-item symmetry and whole-array symmetry. In within-item studies, individual items in the search array either do or do not have internal symmetry, and participants are asked to either find a symmetrical target among asymmetrical distractors, or an asymmetrical target among symmetrical distractors. This approach has provided some evidence of parallel processing of within-item symmetry (Javadnia & Ruddock, 1988), while later work with more diverse and well-controlled stimuli suggested that symmetry detection is a more serial process that requires attention (Olivers and Van Der Helm, 1998). Studies that use the whole-array approach to symmetry in visual search use targets and distractors which do not differ in within-item symmetry but are arranged such that they either do or do not form symmetrical textures across multiple array items. The first to do this was Wolfe and Friedman-Hill (1992) who used oriented lines that were arranged to form symmetrical textures across the search array. Participants were asked to find a target which was oriented such that it disrupted the symmetry of the array. They found that when the distractor arrays were arranged in terms of vertical (mirror) symmetry, finding the target was more efficient than when distractor arrays were arranged in terms of oblique (rotation) symmetry (Wolfe & Friedman-Hill, 1992). Symmetries across the whole array have also been investigated in the context of inter-item symmetries between the target and distractors, and here the findings indicate that when reflection symmetry exists between the target and the distractor, search efficiency is diminished (Zoest et al., 2006). The effect was stronger for vertical than horizontal reflection and is likely the result of shapes that are identical when reflected over the vertical axis being considered as more similar (Zoest et al., 2006). The current study takes inspiration from both approaches and uses a design which allow us to manipulate both within-item and whole-array symmetry in a highly controlled manner.

We used the visual search task to address two research questions: First, how within-item symmetry type (reflection vs. rotation) influences efficiency of visual processing; Second, how texture regularity, a whole-array property, influences the efficiency of visual processing. We contrasted two wallpaper groups: PMM (Figure 2B), which contains 4-fold (90°) reflection and 2-fold (180°) rotation centered at the intersection of the reflection axes; and P4 (Figure 2A), which contains 4-fold (90°) and 2-fold rotation, but no reflection. Both groups (and in fact all wallpaper groups) are textures in which a lattice is repeated to tile the image plane, and the groups differ only in terms of the symmetries within the lattice. The target stimulus was a random dot pattern which contains no internal symmetry and replaced one of the repeating lattices when target was present. The choice of using PMM and P4 as stimuli makes it possible to easily generate exemplars that belong to one group or the other through a very simple image-level operation (see Figure 4 and the Stimuli section of the Methods), and thus manipulate symmetry while controlling for every other image-level attribute (e.g., spatial frequency, contrast, shape of repeating region). Our manipulation of wallpaper group allows us to investigate the effect of within-item symmetry. In order to investigate the effect of regularity across the arrays, we developed “jitter stimuli”, in which gaps were introduced between the repeating lattices, and lattice positions were jittered, such that the regularity of the textures was disrupted (see Figure 2). Regularity is by definition a whole-array property and is therefore related to manipulations of whole-array symmetry in previous studies. This means that our design makes it possible to separately measure effects of within-item symmetry and whole-array regularity. As for the non-jittered displays, the target stimulus was, again, a random dot pattern that replaces one of the array lattices. It is worth noting that across all of our experiments, the visual search task is effectively a search for the absence of symmetry. We will discuss the implications of this in more detail in our General Discussion. Our manipulations of the two dimensions of interest: symmetry type (PMM vs. P4) and texture regularity (no jitter vs. jitter between lattices), give rise to a 2 ✕ 2 design across four visual search experiments, with four array sizes per experiment.

**Figure 2:**
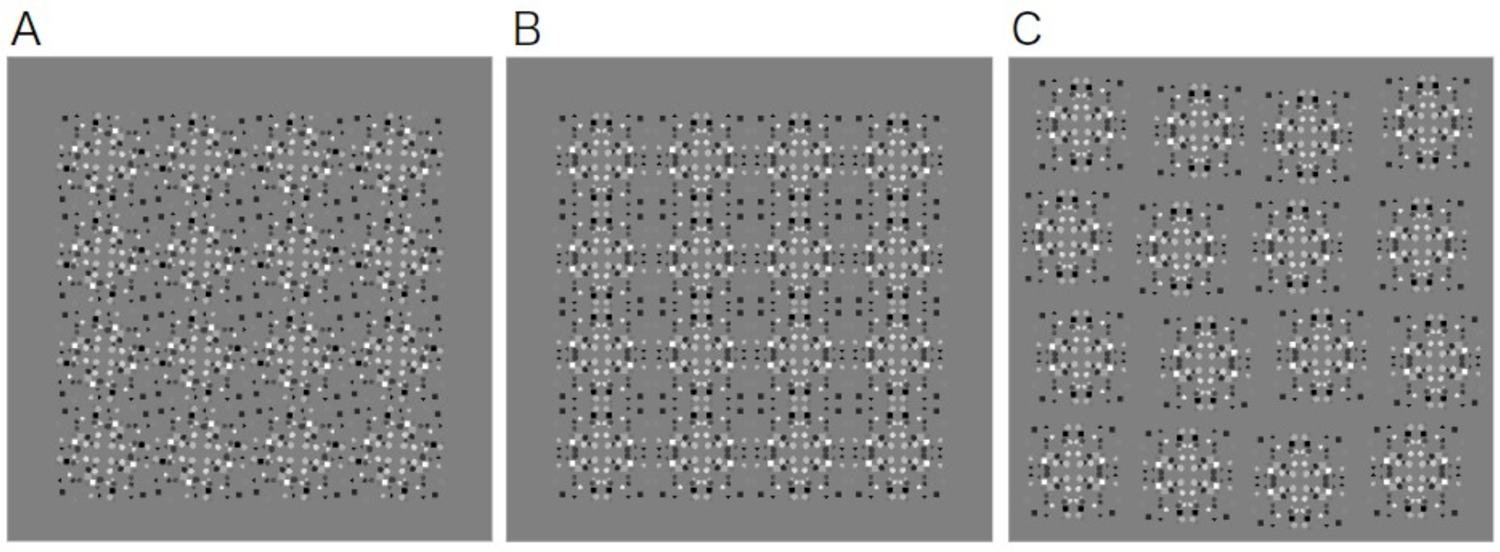
**A**. An example of a P4 wallpaper stimuli, **B**. An example of a PMM wallpaper stimuli. **C**. An example of a PMM “jitter stimuli”. Across all stimuli the individual tiles were the same size and shown on a 50% grey background so the wallpaper would be seamless on the edge. In the jitter stimuli, the overall array is larger, but the tile size is the same.

We used the slope of the linear search function to describe how reaction time and sensitivity (d’) change with larger search arrays as a measure of processing efficiency. Steeper search function slopes indicate that processing is more serial, while shallower slopes indicate more parallel processing, with perfectly flat search function indicating fully parallel processing.

Based on previous results indicating that reflection is processed more efficiently than other types of symmetry, our first hypothesis was that we would find more parallel processing for reflection than for rotation. Our second hypothesis was that regular textures would be processed more efficiently than non-regular textures, as the target would disrupt regularity and perhaps lead to a form of pop-out effect. Our results support the first hypothesis: Across both jittered and un-jittered conditions, reflection symmetries produced more parallel processing. We also confirmed our second hypothesis: search was more efficient for regular textures across both types of symmetry. There were no interactions between symmetry type and regularity, suggesting that the effect of regularity was independent of the effect of symmetry type. These findings add new evidence to the literature on differential processing of reflection and other types of symmetry and demonstrate a novel processing advantage for regular textures.

## 2. Methods

### 2.1 Stimuli

The stimuli were square arrays of lattices. Each lattice was created based on a random dot pattern called a fundamental region, repeated into a 2 x 2 matrix. Two different sets of transformations were applied to the fundamental region inside the lattice. In PMM lattices, the fundamental region is reflected along the vertical axis and then again reflected along the horizontal axis. This produces reflection symmetry along both axes. In P4 lattices, the fundamental region is rotated 90 degrees clockwise, starting with 0 degrees in the upper left quadrant, then 90 degrees to the upper right, 180 degrees in the lower right, and 270 degrees to the lower left. This produces a 4-fold rotation centered at the center of the lattice (see Figure 3). Importantly, all four experiments used the same random dot patterns as fundamental regions, meaning that image-level properties were matched across conditions. When lattices are used to tile the plane, they form regular textures known as wallpaper groups – PMM lattices produce wallpaper group PMM, and P4 lattices produce group P4. We used 10 different fundamental regions to create 10 exemplars of each wallpaper type.

**Figure 3:**
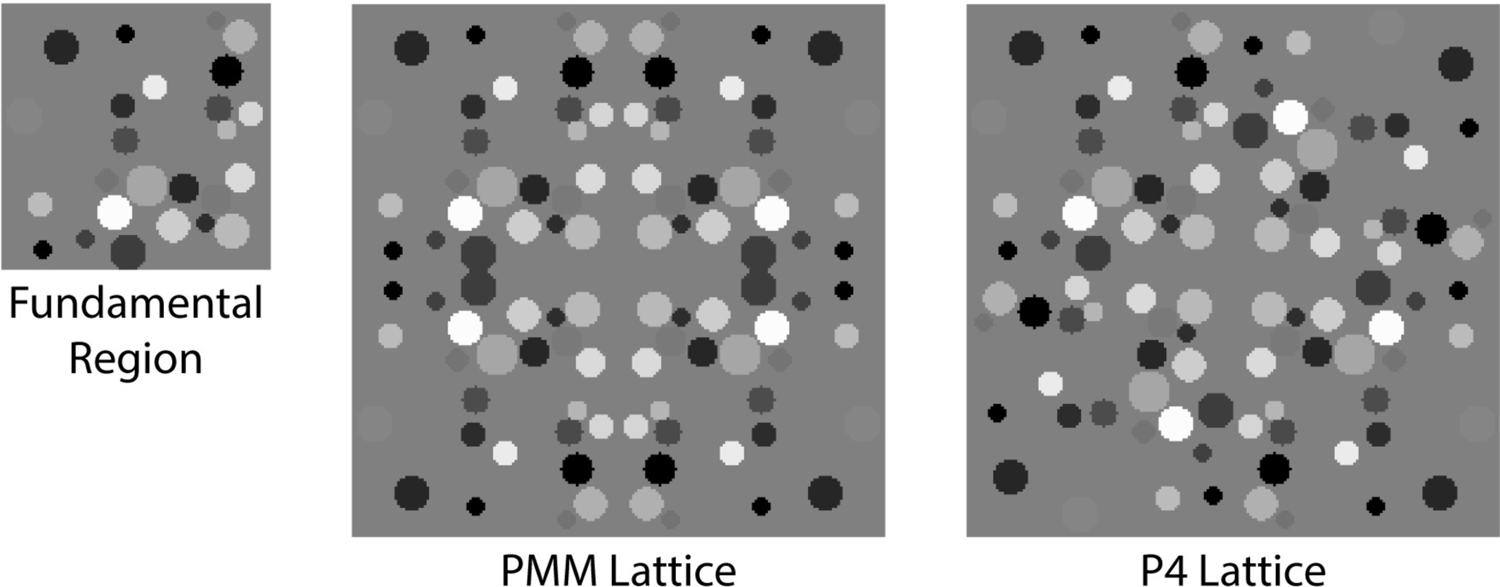
An example of a fundamental region and corresponding PMM and P4 tiles. Here, you can see that the random dot pattern from the fundamental region is repeated 4 times to create a tile but using different transformations. This allows for consistency between the two symmetry types regarding the amount of white, black, and shades of grey in each stimulus.

We also created a “random lattice” that contained no symmetry by using four distinct fundamental regions in each quadrant of the 2 x 2 matrix. These random lattices can be embedded in the wallpaper group in place of any PMM or P4 lattices and are matched to the symmetry lattices in terms of number of dots, contrast and spatial frequency. In our visual search task, the random lattice serves as the target and wallpaper group lattices serve as distractors (see Figure 4). As noted in the Introduction, this participants’ ability to identify the absence of symmetry in the search arrays is used as a measure of symmetry processing, across different conditions. We manipulated symmetry type by either using PMM (“reflection”) or P4 (“rotation”) textures. We further manipulated regularity by adding a separate set of conditions were spacing was introduced around each lattice corresponding to 20% of the tile width/height, and the position of each lattice was jittered randomly between ±15% in both the x and y direction. The jittered conditions were contrasted with un-jittered conditions were there was no gaps between the lattices, so that the distractors formed regular textures. Across all experiments, the lattices were arranged in 3 × 3, 4 × 4, 5 × 5, and 6 × 6 wallpapers to create different array sizes. The size of the lattice was 100 by 100 pixels across all experiments.

**Figure 4:**
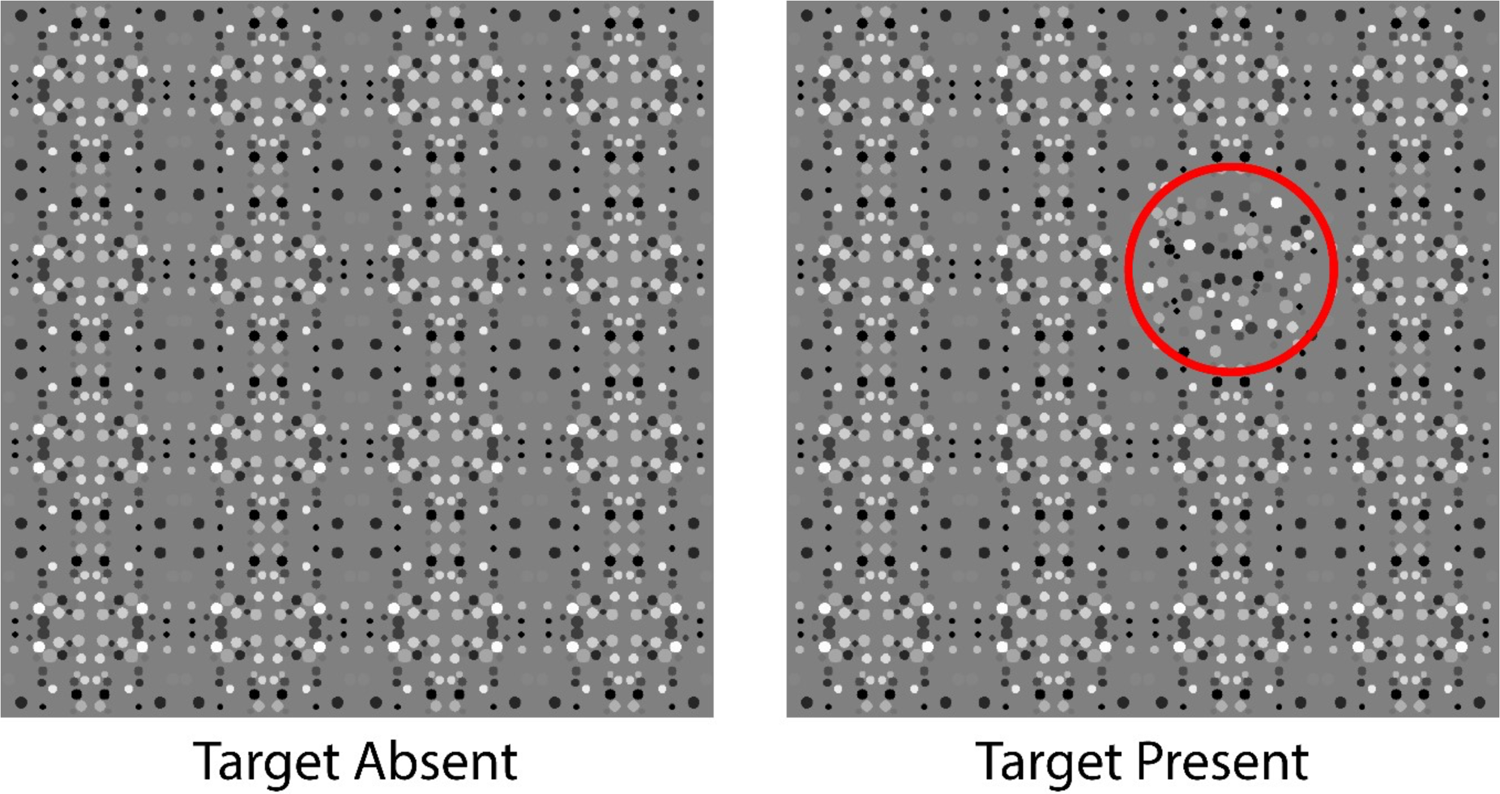
Example images from experiment 1; PMM target absent vs. present example. Circled in red is the “random lattice” target tile.

### 2.2 Participants

Participants were recruited through the online participant pool, Prolific. They were compensated £8.50/hr for their participation, and the experiment lasted about 20mins. For each of our four experiments, we collected data from 50 participants, for a total of 200 (Males = 115, Females = 85). The average age of all participants was 22.46±3.21. Across experiments, we removed participants that were apparently unable or unwilling to do the task (Experiment 1 (PMM) = 6; Experiment 2 (P4) = 10, Experiment 3 (PMM jitter) = 6, Experiment 4 (P4 jitter) = 12), based on a criterion explained below. Informed consent was obtained before the experiment under a protocol approved by the Office of Research Ethics at York University.

### 2.3 Procedure

All four experiments were written using JsPsych (de Leeuw, 2015), hosted online on Pavlovia.org, and followed the same general procedure. Participants were presented with one block of 24 practice trials, followed by 240 experimental trials broken into 10 blocks. The wallpaper array contained a target tile on 75% of trials. Fundamental region exemplars were pseudo-randomly assigned to target-present and target-absent trials so that each of the 10 exemplars was repeated an approximately equal number of times for target-absent and target-present trials, across all array sizes. Trials were shown in random order across exemplars, trial types, and array sizes. When targets were present, their location in the search array was chosen randomly on each trial. Participants were asked to use their keyboard to indicate if a target tile was present or not, pressing the “L” key to indicate that the target was present, and the “D” key to indicate that it was not. Trials only progressed after a selection had been made. After both practice and experiment trials, feedback was provided in the form of the word “Correct!” in green text or “Incorrect!” in red text, presented with a statement indicating how many trials the participant remained in the experiment. Feedback remained on the screen for 1000 ms before the next trial was presented. After each of the 10 blocks, the participants were given the opportunity to take a break before pressing any key to continue. At the end of the experiment, participants were thanked for their participation and redirected to a page where payment could be assigned.

### 2.4 Data analysis

We followed the procedure for calculating *d’* outlined by Macmillian and Kaplan (1985). When individual participants had hit and/or false alarm rates that were 1 or 0, we corrected by adding or subtracting half a trial:

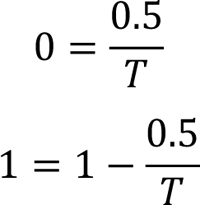

where *T* is the total number of target-present (if correcting hit rates) or target-absent (if correcting false alarm rates) trials. This allowed us to use the standard *z*-score distribution. Participants who had *d’* < 1 were considered unable or unwilling to do the task and removed from further analysis. All statistical analyses was done using JASP (Version 0.18.3).

## 3. Results

We computed the median reaction time and *d’* for each array size, for each participant in each of the four experiments. To test our two hypotheses, we ran a linear mixed models analysis (LMM) separately on the reaction time and *d’* data. Symmetry type (wallpaper group) and jitter were between-subject fixed effects, array size, treated as a continuous variable, was a within-participant fixed effect, and the participant was a random effect. For illustration purposes, we also computed the slope of the search function for reaction time and *d’* individually for each participant (averages across participants are shown in Figures 5B and 6B). Greater slope values are indicative of more serial processing.

**Figure 5:**
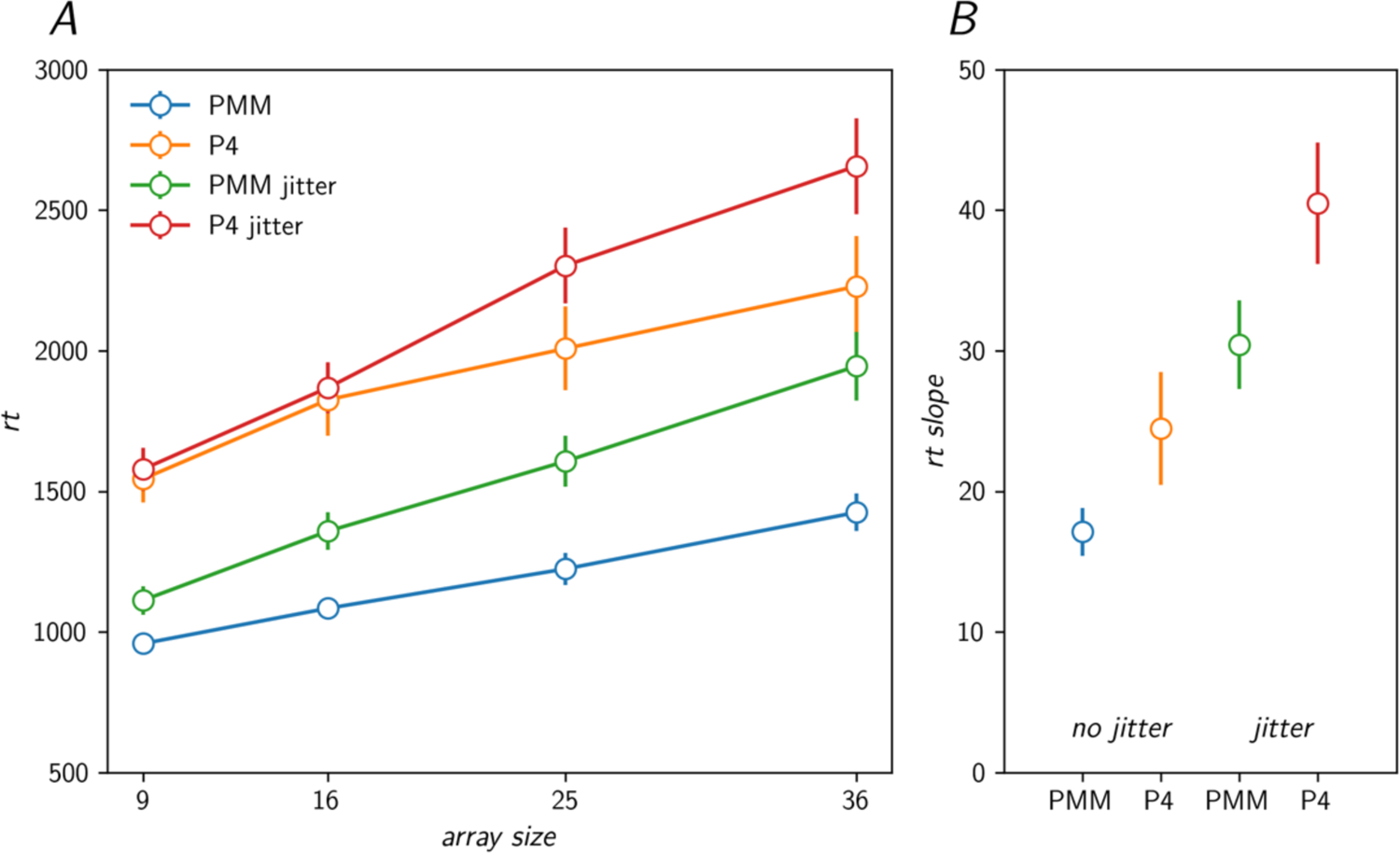
A. Reaction time data across the four experiments. Error bars reflect standard error of the mean. B. Slopes of the visual search function, averaged across participants, for each of the four experiments. Error bars reflect standard error of the mean. It is evident that slope values are smaller (more parallel) for PMM than for P4, and for non-jittered compared to jittered conditions.

**Figure 6:**
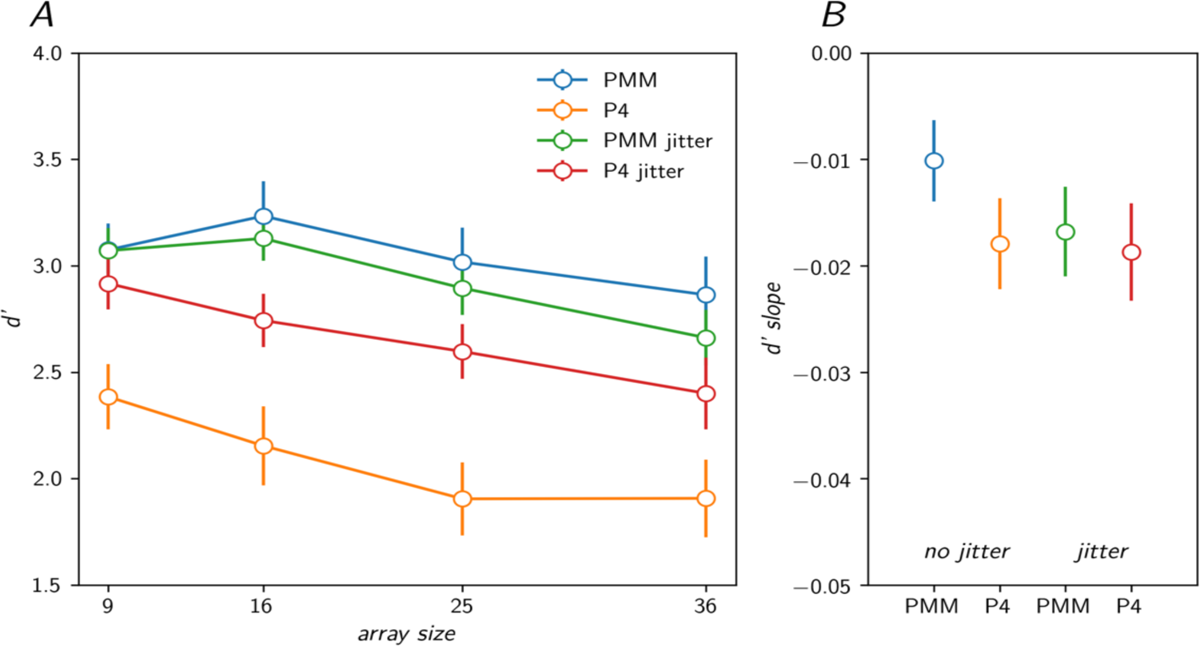
*d’* plotted in the same way as reaction time. Slopes across array sizes are relatively flat, and similar across conditions. The only exception is un-jittered PMM, where the slope is flatter than the others. This implies that this condition was the easiest overall, which is consistent with the reaction time results.

For reaction time, we found a significant main effect of symmetry type (*F*(1,152) = 83.736, p < 0.001), indicating that participants were faster for reflection symmetry (PMM) than for rotation symmetry (P4) across all array sizes. There was also a significant main effect of array size (*F*(1,152) = 283.198, p < 0.001), indicating that reaction time increased with larger array sizes across conditions. Importantly, the significant main effects were modified by two significant interactions that elucidate the efficiency of the visual search: The first interaction was between symmetry type and array size (*F*(1,152) = 6.776, p = 0.010) such that rotation symmetry (P4) produced steeper search functions and thus less efficient search than reflection (PMM). The second interaction was between jitter and array size (*F*(1,152) = 19.258, p < 0.001), such that jittered conditions produced steeper search functions and thus less efficient search than un-jittered conditions. There was no main effect of jitter (p = 0.345) but the interaction between symmetry type and jitter approached significance (*F*(1,152) = 8.403, p = 0.068). Importantly, we did not find a three-way interaction (symmetry type ✕ jitter ✕ array size) (*F*(1, 152) = 0.165, p = 0.685), suggesting that symmetry type and jitter have separate and independent effects on processing efficiency.

We ran the same analysis with *d’* values as the dependent variable to determine if sensitivity was influenced by symmetry type and jitter, and to check for potential speed accuracy trade-offs. As for the reaction time analysis, we found significant main effects of symmetry type (*F*(1,152) = 14.554, p < 0.001), jitter (*F*(1,152) = 5.470, p = 0.021), and array size (*F*(1,152) = 55.029, p < 0.001). The significant main effects were modified by a significant interaction between symmetry type and jitter (*F*(1,152) = 4.317, p = 0.039), but there were no other main effects or interactions (smallest p = 0.258). The slopes were relatively flat and similar across conditions. The only exception is un-jittered reflection symmetries (PMM), which had a flatter slope than the other conditions (see Figure 6). This is consistent with the reaction time analysis, which shows that out of all the conditions, un-jittered reflection symmetries (PMM) led to the most efficient processing.

## 4. Discussion

Our results identify independent effects of symmetry type and texture regularity on visual search efficiency. This is captured by the interactions between symmetry type and array size, and between jitter and array size, that we observe for reaction time. The interactions show that symmetry type and jitter both influence the slope of the search function, with reflection leading to shallower slopes than rotation, and un-jittered displays leading to shallower slopes than jittered displays. These results can be observed in Figure 5, where the shallowest slope is observed for un-jittered reflection symmetry (PMM) and the steepest for jittered rotation (P4). The absence of a three-way interaction indicates that symmetry type and jitter have separate and independent effects on processing efficiency. The pattern of results for *d’* allows us to rule out speed-accuracy trade-off as an explanation for our reaction results.

### 4.1 Behavioral literature on symmetry, reflection vs rotation

Previous research generally used symmetry detection tasks when comparing reflection versus rotation symmetries and found that reflection was more perceptually salient than rotation (Mach, 1959; Royer, 1981; Palmer, 1991; Ogden et al., 2016; Hamada and Ishihara, 1988). We extend these previous findings by demonstrating an advantage for reflection in the context of visual search. We can speculate that the advantage of reflection may be a result of evolutionary pressures, because reflection contributes to the identification of bilateral organisms: members of the same species, predators, and prey. The rapid detection of reflection symmetries would thus be pertinent to survival in hunting, fighting or mating scenarios. It is easy to imagine that these pressures would apply to rotation symmetry detection to a lesser degree, although rotation symmetry may still facilitate identification of various plants, insects, marine animals.

Our texture regularity manipulation reveals a novel processing advantage for symmetries when presented in regular textures. This finding is similar to previous demonstrations of whole-array effects of symmetry in visual search, but in our case, we are manipulating regularity rather than symmetry. The ecological relevance of this effect may be related to evolutionary pressures towards detecting disruptions in regular and near-regular textures in the environment, in the context of detecting edible plants, predators or prey, that are embedded in the background vegetation.

### 4.2 Visual Search and Symmetry

Studies of symmetry using visual search has identified effects of both within-item symmetry (Javadnia and Ruddock, 1988; Olivers and Van Der Helm, 1998) and symmetry over the whole array (Wolfe and Friedman-Hill, 1992). The current study builds on previous work in two important ways: First, by making it possible to independently manipulate symmetry type, as a within-item manipulation, and regularity, by definition a whole-array manipulation. Second, to our knowledge, no prior research has examined how rotation symmetry, whether within a target or a distractor, influences performance on a visual search task. Our study allowed us to place each of our conditions along a spectrum of parallel and serial processing and measure the effect of symmetry type and regularity on both.

We did not see evidence of parallel processing for any of our conditions, while previous work using both within-item (Javadnia and Ruddock, 1988) and whole-array manipulations (Wolfe and Friedman-Hill, 1992) found evidence of parallel processing of reflection symmetry. We believe there a few reasons why that may be. First, while our regularity manipulation is a whole-array manipulation in the same class as that used in previous work demonstrating parallel processing (Wolfe and Friedman-Hill, 1992), we are manipulating regularity while previous work manipulated symmetry. When compared to the previous findings, our results thus suggest that whole-array symmetry, but not whole-array regularity, give rise to parallel processing.

Our within-item manipulation of symmetry, however, is more generally similar to the types of displays used in previous work (Javadnia and Ruddock, 1988; Olivers and Van Der Helm, 1998), especially in our jittered conditions. Should we be surprised to not see parallel processing when within-item reflection or rotation symmetry differed between the target and the distractor? To our knowledge, the evidence of parallel processing in within-item symmetry-based visual search is limited to a study (Javadnia and Ruddock, 1988) that used stimuli based on “textons” (Julesz, 1981). Here, the target and distractor stimuli did differ on symmetry, but also differed in other respects, such as the presence of line junctions of various types, which may have facilitated pop-out of the target among the distractors. A more recent study used more well-controlled stimuli and found no evidence of parallel processing of symmetry across four distinct stimulus types (Olivers and Van Der Helm, 1998). The stimuli used in the current work has targets and distractors that are similarly well-matched for low-level differences, and further allows us to manipulate symmetry type without introducing any low-level differences. Given this high level of control, we are not surprised that we, like Oliver and Van Der Helms (1998), find no evidence of parallel processing of within-item symmetry.

### 4.3 Neuroimaging literature on symmetry, reflection vs rotation

The neuroimaging literature shows that both reflection and rotation produce strong responses in the visual cortex, even when participants are doing an orthogonal task and not paying attention to the symmetry (Kohler et al., 2016), but activity measured using EEG was weaker for rotation than for reflection symmetry (Kohler & Clarke, 2021). A recent direct comparison of responses to different wallpaper groups in the visual cortex of macaque monkeys showed that activation in visual areas (V3 and V4) was approximately 1/3 larger for reflection than for rotation (Audurier et al., 2021). Studies using non-texture stimuli with a single symmetry axis have also consistently found weaker responses for rotation than reflection (Makin et al., 2012, 2013, 2014; Wright et al., 2015). This neural advantage for reflection over rotation is consistent with the behavioral advantage observed in the current study and prior studies discussed above. An important goal for further neuroimaging research will be to directly compare symmetries when presented independently or embedded in regular textures to understand how the behavioral advantage for symmetries in regular textures arises in the brain.

### 4.4 Texture perception

Textures form the patterns that make up the surfaces of objects and environments; they play an essential role in vision in everyday life (Adelson, 2001). An important step toward understanding and analyzing human texture perception was the development of a computational model that made it possible to represent and synthesize visual textures based on joint statistics of the image (Portilla & Simoncelli, 2000). The model has proven highly useful in capturing how texture representations change across the visual field (Balas, Nakano, and Rosenholtz, 2009; Freeman & Simoncelli, 2011) and how natural textures are represented in different areas of the visual cortex (Freeman et al., 2013; Okazawa et al., 2015). Importantly, however, this modeling framework is unable to synthesize regular textures like the wallpaper groups used in our experiment (unpublished data) and therefore unlikely to fully explain the regularity effect found in the current data or the brain imaging data mentioned above. The current data offer another piece of evidence suggesting that regular and near-regular textures may play an important role in perception. Our findings highlight the need for the development of models that can describe and synthesize regular textures.

The uniformity illusion (Fukuda & Reno, 2012; Otten et al., 2016) may also be relevant to our results. In this illusion, a regular grid of repeating local elements can undergo illusory completion such that a disruption of the uniformity in the periphery is not detected. The illusion suggests that the visual system has a tendency towards perceiving textures as regular, which implies that visual search should be less efficient when the target is presented in a regular texture, because the target is filled-in to give the impression of a uniform pattern. Why do we see the opposite pattern, more efficient search for regular textures? One possibility is that the mechanism underlying the uniformity illusion only operates in the far periphery, far enough to not influence our results. Another, related, possibility is that the uniformity illusion requires more tiling of the repeating pattern, than is present in our textures. Otten and colleagues (2016) used relatively large central segments (e.g., 26 x 14 °/visual angle), and while they did not systematically test the effect of tiling, the number of repetitions did in most cases exceed that present in even our largest stimuli. It is worth noting that while these differences could potentially explain the absence of a regularity effect in our results, they do not predict the effect we see in the opposite direction. It is compelling to consider a third possibility, namely that the filling-in mechanism is unable to reproduce the local symmetries, whether reflection or rotation, present in our stimuli. If the visual system has a tendency towards perceiving textures as regular, it is plausible that this failure to fill-in the regularity would give rise to a strong error signal, that would in turn make visual search more efficient with a regular search array. Otten and colleagues (2016) have demonstrated that the uniformity illusion can be produced for a wide range of features, and the wallpaper group stimuli used in the current study provide an ideal stimulus set for adding symmetry to that list. Further work should investigate which parameters are required for generating a potential uniformity illusion with repeating elements that have local symmetries.

### 4.5 Potential confounds and limitations

A possible limitation of this study is that while the stimuli were well controlled within participant, the data were collected online on participants own devices, which has the potential for introducing differences between participants. Participants were required to use a laptop or desktop computer for the experiment (no phones or tablets were allowed), but we made no attempts to control viewing distance or monitor resolution, which likely lead to some variability in the size of the stimuli in degrees of visual angle between participants. In addition, the contrast and luminance of the stimuli may also have varied because of differences in the monitors’ used by different participants. It is important to note, however, that our effects of interest were measured within-participant, and thus unlikely to be driven by these between-participant differences. Furthermore, any noise added to our measurements by the lack of control is likely compensated for by our ability to get data from a relatively large number of participants, compared to a standard psychophysical experiment.

A second possible limitation is that the target was always asymmetrical, while the distractors were symmetrical. We deliberately chose this approach because it allows us to organize the search arrays into wallpaper groups PMM and P4, thereby connection our results to the existing literature on visual processing of symmetries in wallpaper groups (e.g., Kohler et al., 2016). We cannot rule out that using a symmetrical target among asymmetrical distractors would have led to pop-out with our stimuli, but from casual observations we consider it highly unlikely. It is also worth noting that previous work used both symmetrical and asymmetrical targets, found evidence of parallel processing within asymmetrical targets, and took that as evidence of parallel processing of symmetry (Javadnia and Ruddock, 1988). Similarly, Wolfe and Friedman-Hall (1998) measured differences in visual search patterns through the disruption of whole-array symmetry (introducing asymmetry to a symmetrical array) rather than the identification of symmetry components within an asymmetrical array. Based on this, we conclude that using an asymmetrical target to measure symmetry processing is reasonable and well-aligned with the existing literature.

Finally, a possible concern regarding our regularity manipulation is that the regular arrangement of the lattice in the wallpaper stimuli may have produced a mid-level visual effect where individual dots in the lattices are perceptually grouped to form a grid-like pattern. It is possible that this can help guide the visual search task, because the target (a disruption of the grid) becomes easier to spot. We have several responses to this concern: First, it is important to note that such patterns across the whole texture are in a sense inherent to regular textures, and thus difficult to disambiguate from regularity itself. Second, we note that the presence of a grid pattern does not necessarily mean that a disruption of the grid becomes easy to detect, as evidenced by the uniformity illusion (discussed above; see Figure 7 in Otten et al., 2016). Finally, if the grid pattern was driving our regularity effect, we would expect there to be a stronger effect of regularity on PMM, where straight lines are more likely to form across the pattern (see Figure 4). This should produce a 3-way interaction between symmetry type, jitter, and array size. However, we did not find any such interaction, so we believe the grid effect is unlikely to impact our results.

## 5. Conclusion

The current study used highly controlled stimuli to provide strong evidence that reflection symmetries (wallpaper group PMM) are processed more efficiently than rotation symmetries (wallpaper group P4). Perhaps more surprisingly, the results also show that texture regularity has a significant effect on the processing of symmetries independent of symmetry type, such that when symmetries are embedded within a regular texture, they are more efficiently processed. Notably, these effects were independent and additive, which suggests that they reflect distinct perceptual mechanisms. As pointed out above, a major distinction of our work from previous work on visual search is our ability to test both within-item and whole-array manipulations in the same display and show that within-item symmetry type as well as whole-array regularity matters for visual search. The findings should prompt further studies of the underlying brain mechanisms and provide a foundation for further research on how symmetries and textures interact during natural, real-world perceptual tasks.

## 6. Data Availability

The datasets collected for the current study are available in the project wallpaper-search repository, https://osf.io/b7hq9/

## Notes

https://osf.io/b7hq9

### Competing Interest Statement

The authors have declared no competing interest.

